# The prevalent *NR2E3* c.932G>A mutation induces aberrant splicing that can be rescued using splice-shifting antisense oligonucleotides

**DOI:** 10.1101/2024.05.01.592034

**Authors:** Yehezkel Sztainberg, Maya David Teitelbaum, Ilana Buchumenski, Hagit Porath, Dror Sharon, Eyal Banin, Rotem Karni, Erez Y. Levanon, Ariel Feiglin

## Abstract

Mutations in *NR2E3* have been implicated in several progressive retinal disease phenotypes such as enhanced S-cone syndrome, Goldmann-Favre syndrome and retinitis pigmentosa. One of the most frequent mutations in NR2E3 is c.932G>A (p.R311Q), where pathogenicity is thought to stem from the resulting amino acid substitution. However, multiple studies that evaluated the effect of this substitution on the protein, did not elucidate the molecular basis underlying the pathogenicity.

Primed by bioinformatic analyses, we hypothesized and experimentally validated that the *NR2E3* c.932G>A mutation leads to aberrant splicing which results in a short, non-functional protein isoform. Using cell models expressing WT and mutant constructs of the full *NR2E3* sequence (including exonic and intronic regions), we observed that the mutated transcript exhibits a high level (75%) of aberrant splicing through gain of a novel splice acceptor site within exon 6. This mis-splicing results in the in-frame loss of 186 base pairs that code for a portion of the protein ligand binding domain. We further designed and evaluated splice-shifting antisense oligonucleotides (ASOs), that circumvented the aberrant splicing. The best performing ASO successfully restored 70% of the total NR2E3 full-length isoform levels and demonstrated rescue of nuclear localization and rhodopsin transcriptional activation.

This study demonstrates the importance of understanding splicing consequences of pathogenic mutations, allowing the design and development of ASO-based therapies. Our findings set the stage for the potential treatment of *NR2E3*-related retinal degeneration caused by the c.932G>A mutation using splice-shifting ASOs.

## Introduction

Inherited retinal diseases (IRDs) represent a diverse group of genetically heterogeneous disorders than can lead to progressive vision impairment and blindness ^1,2^. Collectively, IRDs are estimated to affect more than 5 million people worldwide ^3^. IRDs are predominantly monogenic diseases, mainly caused by mutations in genes that are critical to retinal function. To date, mutations in more than 270 genes have been identified to cause IRDs ^4^.

The *NR2E3* (nuclear receptor subfamily 2, group E, member 3) gene encodes an orphan nuclear receptor that functions as a ligand-dependent transcription factor essential for the normal development and differentiation of rod photoreceptors ^5,6^. Mutations in *NR2E3* were initially described in patients with enhanced S-cone syndrome (ESCS) ^7^, but later were associated with other inherited degenerative retinal phenotypes including Goldmann-Favre syndrome (GFS), clumped pigmentary retinal degeneration (CPRD), and retinitis pigmentosa (RP) ^8^. ESCS is an autosomal recessive retinopathy characterized by lack of rod photoreceptors and an abnormal increase in the number of blue (S) cones, resulting in night blindness, retinal dystrophy, and gradual loss of vision ^9^.

One of the most frequent pathogenic variants in *NR2E3* is the so-called missense mutation c.932G>A (p.R311Q) located in exon 6 ^7,10–12^. Although this mutation is positioned in the NR2E3 ligand binding domain, several studies have failed to understand its exact mode of action, as the p.R311Q mutant protein in these studies appears to conserve normal DNA binding ^13–15^, homodimerization ^13,14^, interaction with CRX ^14,16^, and transcriptional activity ^14–16^. To the best of our knowledge, only one study, found the p.R311Q mutant to be defective in transcriptional repression of M-opsin, but only under specific conditions out of many that were evaluated ^13^.

These findings, in combination with bioinformatic analyses that indicated a possible splicing effect for this mutation led us to hypothesize that the pathogenicity of c.932G>A stems mainly from the change in splicing rather than the amino acid substitution. In this study we demonstrate that indeed this mutation creates a new splice acceptor site within exon 6 of *NR2E3*, causing in-frame aberrant splicing that translates to the loss of a portion of the ligand binding domain. Based on these results, we developed antisense oligonucleotides (ASO) that correct this aberrant splicing *in-vitro* and successfully restore molecular function of the protein.

## Results

Figure 1 illustrates the bioinformatic prediction and experimental validation of aberrant splicing caused by the *NR2E3-* c.932G>A mutation. We performed a bioinformatic analysis of the *NR2E3* c.932G>A mutation using SpliceAI, an algorithm specialized for identification of splicing signals caused by genomic alteration^17^. SpliceAI predicted with high confidence (score = 0.74, see Methods), the creation of a new splice acceptor site consensus sequence, 186 bp downstream of the original acceptor site. This change is expected to result in loss of the first 186 base pairs of exon 6 (Fig. 1a), known to code for part of the ligand binding domain. In contrast, REVEL^18^ and AlphaMissense ^19^, bioinformatic tools that focus on predicting pathogenicity caused directly by the amino acid change and do not take splicing into account, indicated no strong effect resulting from the Arginine 311 to Glutamine alteration (see Methods). To confirm the splicing prediction, two plasmids expressing the *NR2E3* WT transcript (NR2E3^WT^) and the mutated transcript (NR2E3^c.932G>A^) of the gene were generated (Fig. 1b). Notably, these plasmids were designed to include not only exonic regions but also all *NR2E3* introns (see Methods, Fig. 1b and Supp. Fig. 1). Splicing of these *NR2E3* constructs was evaluated in HEK293 cells. RT-PCR using primers spanning exons 5-7 (see Methods) confirmed the bioinformatic predictions, demonstrating expression of an aberrant, shorter, transcript in the cells transfected with mutant but not the WT plasmid (Fig. 1c). This was further confirmed at the protein level by western blot analysis (Fig. 1d). Sequencing of the PCR product revealed the loss of 186 bases of exon 6, precisely as predicted (Fig. 1e).

**Fig. 1.**
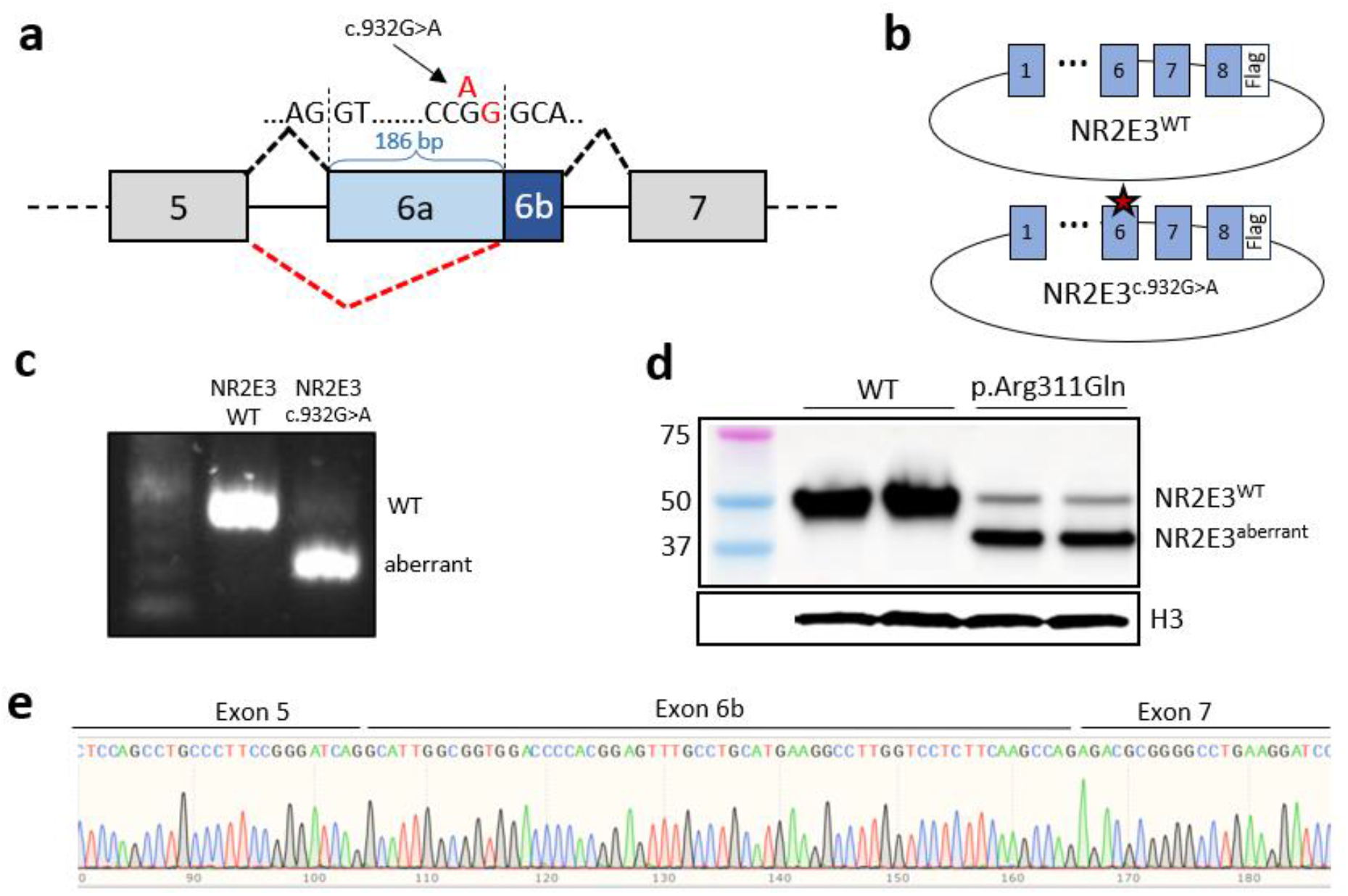
Aberrant splicing of *NR2E3* due to c.932G>A. **(a)** The *NR2E3* c.932G>A mutation is computationally predicted to cause cryptic splicing, resulting in the loss of 75% of exon 6. Part of the genomic sequence is presented above the gene model. The new splice acceptor “AG” site is marked in red characters. **(b)** Design of NR2E3^WT^ and NR2E3^c.932G>A^ plasmids (flag-tagged) with all exons and introns. **(c)** Gel electrophoresis of RT-PCR products obtained from HEK293 cells transfected with NR2E3^WT^ or NR2E3^c.932G>A^ plasmids. RT-PCR was performed using primers spanning exons 5, 6, and 7. **(d)** Western blot analysis confirms aberrant splicing at the protein level. Histone 3 (H3) is used as a loading control. **(e)** Sequencing of the lower band of the RT-PCR product of NR2E3^c.932G>A^ in (c) confirms the predicted aberrant splicing.

A potential therapeutic approach to circumvent the aberrant splicing caused by the *NR2E3-* c.932G>A mutation and thus restore normal splicing, is through the use of splice shifting ASOs. To this end, we designed and evaluated 16 ASOs, spanning 50 bp and flanking the position of the aberrant splice acceptor site in exon 6 (Fig. 2a). A semi-quantitative PCR mini screen was performed following transfection of the mutant plasmid into HEK293 cells and treatment with the different ASOs. ASO-2 and ASO-3 partially corrected the aberrant splicing and restored the expression of the full-length transcript (Fig. 2b). ASO-2 and ASO-3 increased the full-length transcript by 2.2- and 2.3-fold over the control, reaching 28% and 30% of the WT expression levels respectively. Restoration of normal splicing was next demonstrated at the protein level. ASO-2 treated cells demonstrated an 8-fold increase (p-value=0.006) in the fraction of full-length protein compared with the mis-spliced isoform, reaching 70% of the original protein level (Fig. 2c). Notably, restoration of the full-length isoform by ASO-2 was more efficient at the protein level compared to the RNA level. This could be explained by increased stability of the corrected, full-length, protein. ASO-2 was therefore selected as the preferred candidate for additional experimentation.

**Fig. 2.**
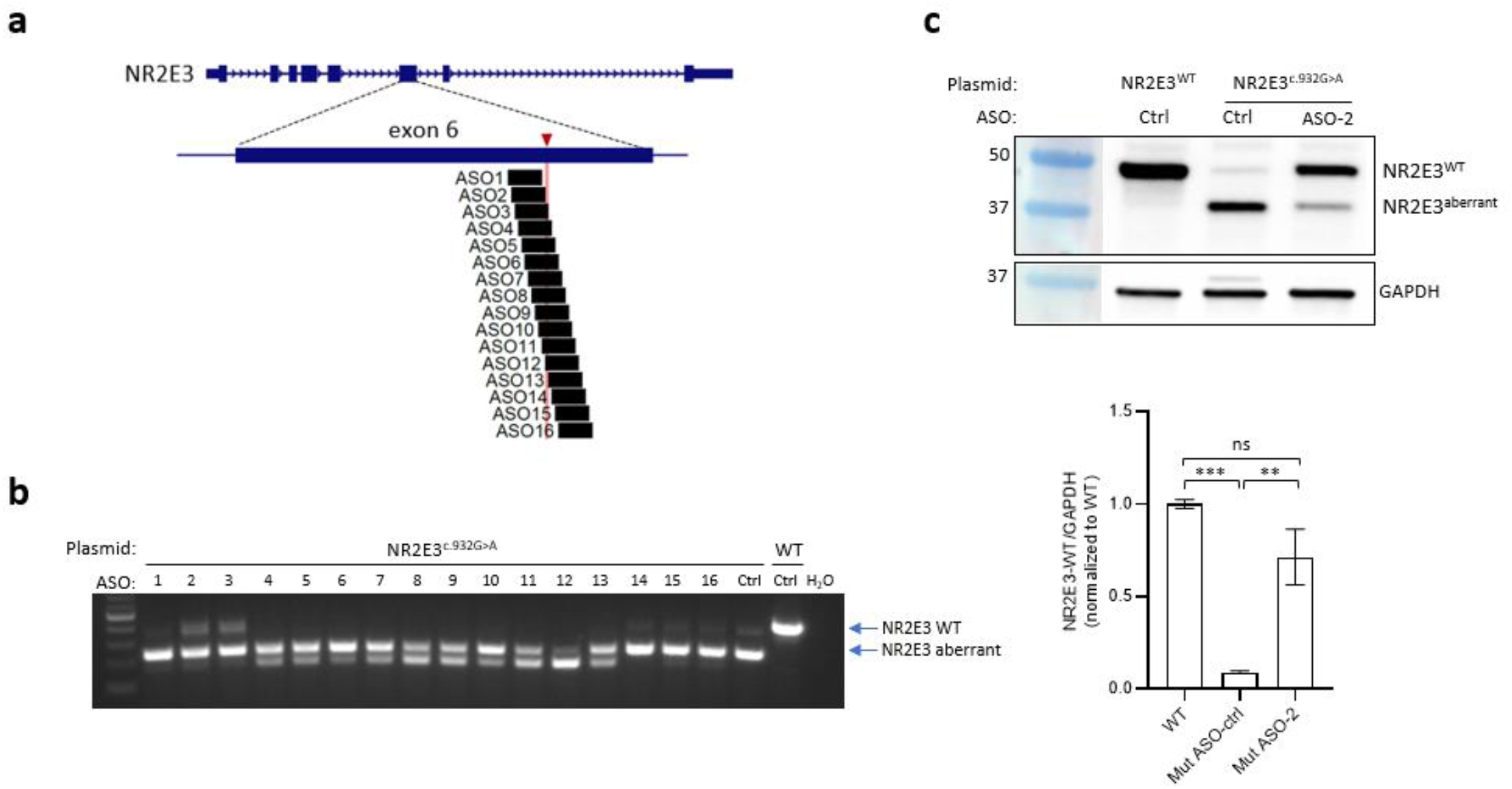
ASOs restore normal splicing in NR2E3^c.932G>A^. **(a)** Snapshot from the UCSC genome browser showing the NR2E3 gene with magnification of exon 6 and mapping of ASOs 1-16. The c.932G>A mutation position is indicated (red triangle and vertical line). **(b)** Oligo-screen with ASOs 1-16 to correct aberrant splicing in HEK293 cells transfected with NR2E3^c.932G>A^ plasmid. RT-PCR was performed using primers spanning exons 5, 6, and 7. Ctrl – antisense control. WT – plasmid expressing NR2E3^WT^. **(c)** Western blot (top panel) and densitometry analysis of the upper band (bottom panel) confirms correction of aberrant splicing at the protein level (average of 3 independent experiments). GAPDH is used as a loading control. Data are mean ± s.e.m. \*\**P*<0.01; \*\*\**P*<0.001 (One-way ANOVA). ns - not significant.

We next evaluated the ability of ASO-2 to restore molecular characteristics of NR2E3 function (Figure 3). NR2E3 is known for its role as a transcription factor and is thus expected to localize in the nucleus of the cell. Therefore, we assessed the nuclear sub-cellular localization of NR2E3 WT and mutant proteins (NR2E3^c.932G>A^) by immunofluorescence following transient transfection of plasmids. We found that whereas NR2E3^WT^ was clearly localized in the nucleus, the mutant, NR2E3^c.932G>A^, was characterized by less intense staining, mostly located outside the nucleus, a characteristic previously reported for several NR2E3 mutations ^20^. Treatment with ASO-2 successfully restored the nuclear localization of NR2E3 (Fig. 3a). Restoration of nuclear localization was further confirmed using nucleus-cytoplasm fractionation. Treatment with ASO-2 increased full-length NR2E3 protein expression in the nucleus 10.6-fold compared to non-treated levels (Fig. 3b and Supp. Fig. 2).

**Fig. 3.**
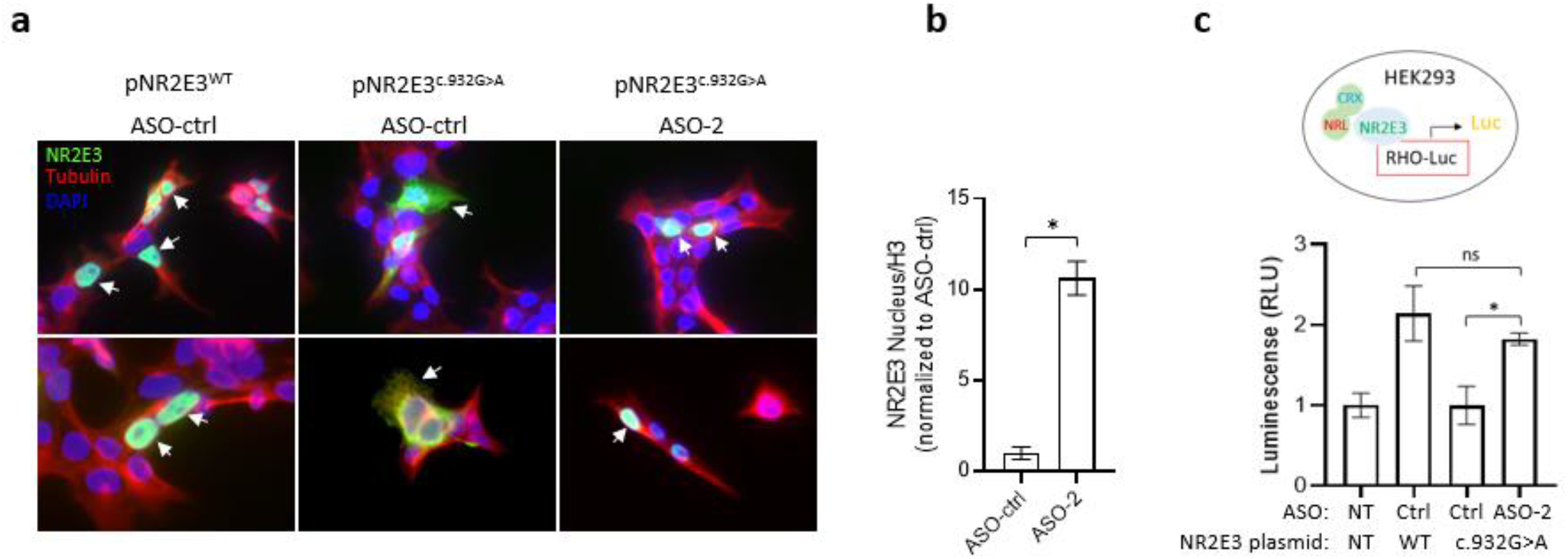
ASO treatment corrects NR2E3^c.932G>A^ sub-cellular localization and RHO promoter activation. **(a)** Immunofluorescence staining following transfection of plasmids and ASO treatment in HeLa cells. Arrowheads indicate localization of NR2E3 expression. **(b)** Densitometry analysis of a nucleus-cytoplasm fractionation experiment in HEK293 cells (see Supp. Fig. 2). **(c)** Luciferase activity assay following ASO treatment (n=4). All 4 groups were transfected with RHO reporter, CRX, and NRL. ns – not significant. Ctrl – ASO control. Data are mean ± s.e.m. \**P*<0.05 (One-way ANOVA).

NR2E3 plays a key role in activation of rhodopsin (RHO) expression in rod cells^21^. Therefore, we further evaluated the ability of ASO-2 to restore this molecular function. To this end, we used a luciferase reporter assay controlled by the rhodopsin promoter in the presence of NR2E3’s known co-factors CRX and NRL^16^. Results show that in this setup, WT NR2E3 induces a 2-fold increase in RHO transcription activation whereas the mutant gene, NR2E3^c.932G>A^, could not induce any transcription. Treatment of mutant cells (expressing NR2E3^c.932G>A^) with ASO-2 resulted in nearly complete rescue of RHO transcription activation, increasing transcription 1.8-fold over the control ASO, and reaching 85% of the WT (Fig. 3c), demonstrating therapeutic potential of ASO-2 treatment.

## Discussion

Pathogenicity of missense mutations is typically thought to stem from the change in the encoded amino acid that either disrupts a functional domain or induces misfolding of the resultant protein. Based on this hypothesis, the classic method to study the effect of a mutant protein *in vitro* is through a vector encoding the mutant cDNA (without introns) and expressing it in a relevant cell line for subsequent phenotypic characterization. However, this approach could lead to an incomplete understanding of the molecular consequences of the mutation as it cannot factor in potential splicing changes triggered by the mutation. These splicing changes can only be revealed when introns are included in the expressing vector.

Bioinformatic tools focused on prediction of splicing changes are gaining traction and are an important tool for highlighting “suspicious” cases. In the case of the *NR2E3* c.932G>A mutation, SpliceAI predicted creation of a new splice acceptor site that leads to the loss of ∼75% (186 of 247 bp) of exon 6. This analysis, coupled with a low bioinformatic score predicted by tools focusing on biochemical changes in the protein, highlighted this mutation as an interesting target to explore. We experimentally confirmed the prediction that the mutation indeed confers aberrant splicing using an expression vector encoding the full-length NR2E3, including all its exons and introns, allowing for the splicing process to occur. Using this vector, we also demonstrated that the protein resulting from the NR2E3 c.932G>A mutation is aberrantly localized in the cytoplasm, suggesting its function as a transcription factor is compromised.

Potential reversibility of phenotype after disease onset, was demonstrated in the rd7 (NR2E3 knockout) mouse model, following treatment with AAV-*NR2E3*. Subretinal injection of the virus rescued the phenotype by protecting photoreceptors from further damage^22^. Testing in additional animal models led to clinical trials in human patients manifesting an RP phenotype caused by NR2E3 and RHO mutations, as well as LCA caused by CEP290 mutations. A phase 3 clinical trial utilizing this AAV5-hNR2E3 vector, called OCU400, was recently approved by the FDA to test its efficacy and safety as a gene-agnostic therapy to modify course of disease in RHO-RP as well as RP caused by other genes ^23,24^. Based on these data, we proposed to use ASOs to correct the splicing defect and restore expression of full-length NR2E3. As c.932G>A is one of the most common NR2E3 mutation, we anticipate a significant number of suitable patients. ASO development is especially relevant for splicing mutations, with successful treatments based on splicing modulation emerging in the clinic ^25–27^. In this context, retinal diseases are especially relevant, as the delivery of ASOs to the various layers of the retina and specifically photoreceptor cells are proving to be efficient. In addition to the use of naked oligonucleotides with various chemical modifications in the retina, the use of delivery vehicles and conjugates specific for retinal delivery are being developed. Therefore, we tested the ability of ASOs to reverse the aberrant splicing caused by the c.932G>A mutation. Evaluating the impact of ASO-based rescue of NR2E3 splicing is a crucial step in the path to therapeutic development. Especially since the ASO-based rescue, in this case, does not restore the WT protein but rather triggers production of the p.R311Q isoform. In this regard, it is important to note that most studies did not observe a distinct phenotype when evaluating this amino acid change^13–16^. Also, the mutation itself does not seem to have a toxic affect, as individuals with only one mutated allele do not develop a clinical phenotype (recessive disease). In this study we were able to demonstrate correction of sub-cellular localization and restoration of rhodopsin transcriptional activation following ASO treatment. These are encouraging first steps that indicate restoration of protein function.

As ASOs are sequence-specific, it is challenging to translate ASOs targeting non-human species to ASO that would be relevant for the human genome. Therefore, an *in-vivo* model, would require a humanized version of the gene carrying the specific mutation. In the absence of such models for most ultra rare diseases and specifically for ESCS, retinal organoids are a logical way forward. Therefore, next steps should include biopsies from patient cells and development of retinal organoids that can be evaluated more extensively than cell lines. This approach will hopefully help advance therapeutic ASO development for rare genetic disorders.

## SUPPLEMENTARY FIGURE 1

**Supp. Fig 1.**
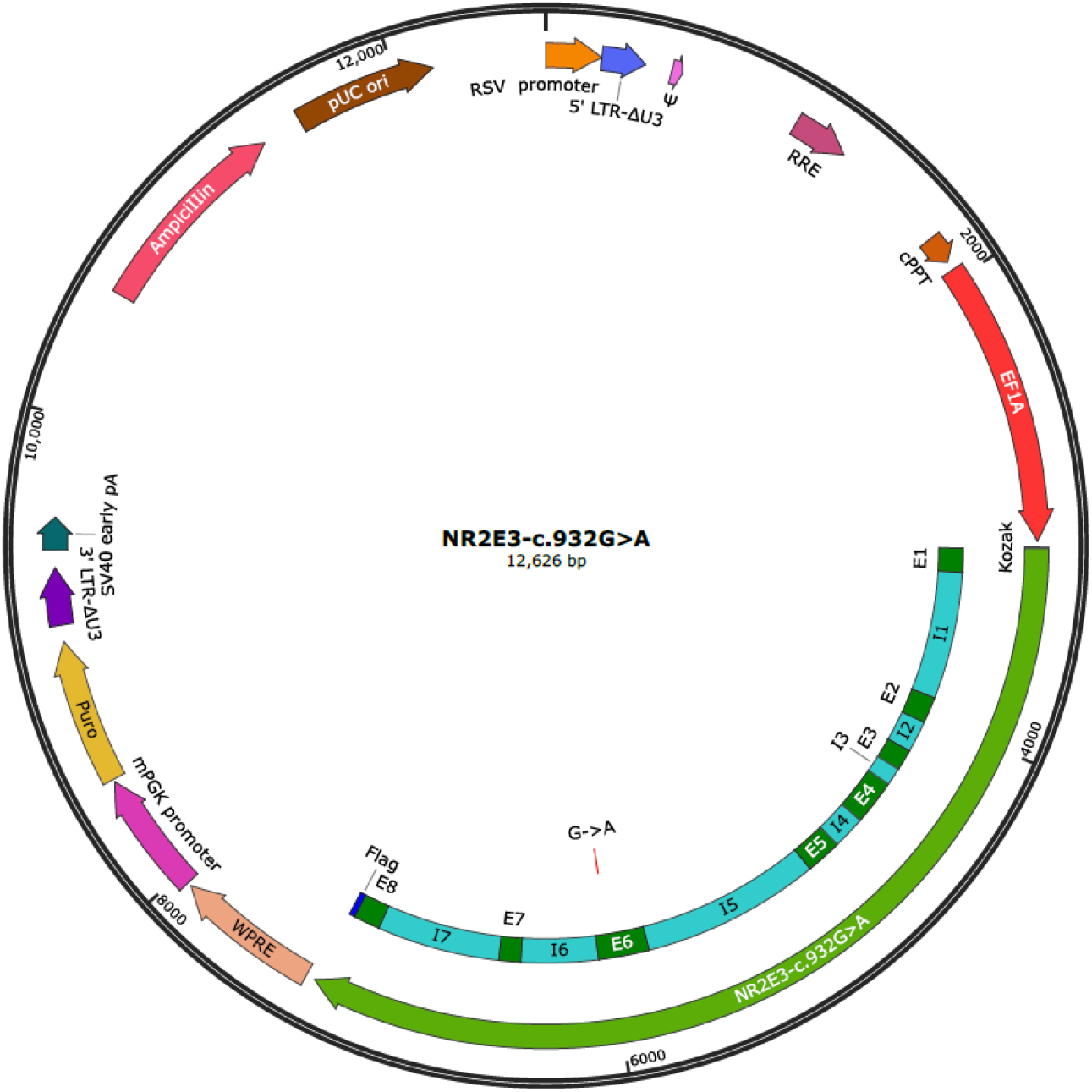
NR2E3 Plasmid design. The plasmids (WT and mutant) include the EF1A promoter, full-length exons and introns of NR2E3 (except for intron 7 that only includes the first and last 300 bp of the original intron sequence), a C-terminal flag tag, and puromycin resistance for selection. In addition, the vectors are suitable for lentiviral packaging.

## SUPPLEMENTARY FIGURE 2

**Supp. Fig 2.**
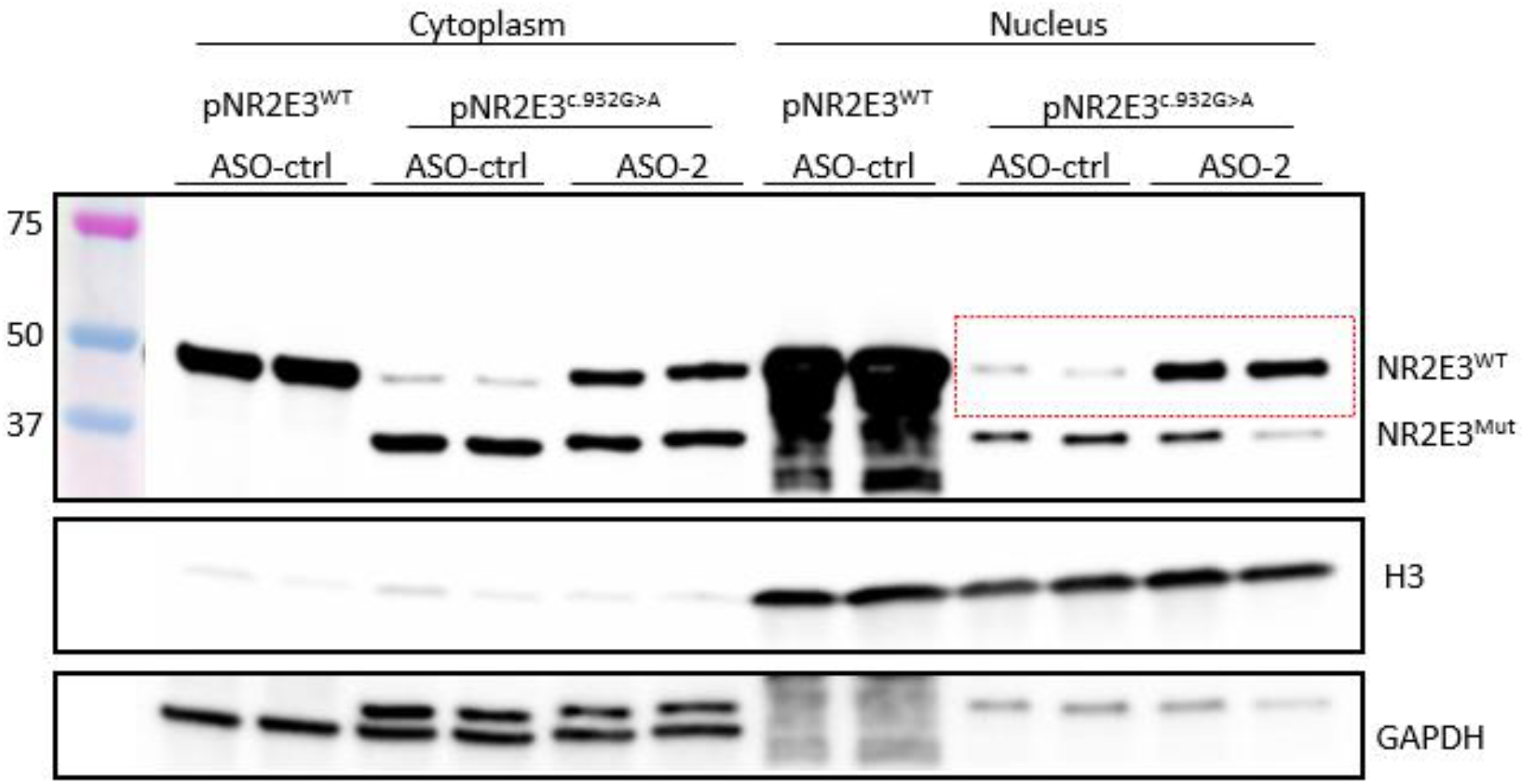
ASO treatment restores NR2E3 expression in the nucleus. Nucleus-cytoplasm fractionation in HEK293 cells transfected with plasmids and treated with ASO. H3 and GAPDH are used to confirm correct nucleus-cytoplasm fractionation. Red/dashed rectangle indicates the bands quantified in Fig. 3b.

## Materials and methods

### Bioinformatic scores

SpliceAI was used through the web interface https://spliceailookup.broadinstitute.org/. Input was G to A substitution at position chr15:71813573 (hg38 genome build). Output was the prediction of a splice acceptor site gain with a score of 0.74 (on a scale of 0-1, with 0 indicating lowest and 1 indicating highest probability for splicing) at position chr15:71813575, 2 bp downstream of the mutation^17^. Revel score of 0.107 (on a scale of 0-1, with 0 indicating benign and 1 indicating pathogenic mutations) was taken from the UCSC browser track: “REVEL Pathogenicity Score for single-base coding mutations” at position chr15:71813573^18^. AlphaMissense^19^ score of 0.093 (on a scale of 0-1, with 0 indicating benign and 1 indicating pathogenic mutations) was taken from supplementary table S5 at Ref. 19 for the chr15_71813573_G_A_hg38 variant.

### Transfection of plasmids and ASOs

NR2E3 plasmids were ordered from VectorBuilder, with C-terminal Flag tag, and EF1a promoter. The plasmids encode the full-length exons and introns of NR2E3, except for intron 7 that only includes the first and last 300 bp of the original intron sequence. HEK293 cells were transfected with 1μg of each plasmid in 6-well plates using FuGene HD transfection reagent, 4 hours after plating the cells. Cells were collected 48hrs after plasmids transfection for either RNA or protein extraction. ASOs were ordered from BioSpring and consisted of 20 RNA nucleotides with full phosphorothioate (PS) and *2*’-O-methoxyethyl (2’-MOE) chemical modifications. For the ASO experiments, HEK293 were first transfected with plasmids on 6-well plates. The next day, cells were trypsinized and replated on 24-well plates. Four hours after replating, cells were transfected with 500nM NR2E3 ASOs or ASO control using FuGene HD transfection reagent. Cells were collected 48hrs after ASOs transfection for either RNA or protein extraction.

### Semi-quantitative PCR

RNA from cells was extracted with the Monarch Total RNA Miniprep (NEB T2010S) following the manufacturer instructions, and 1μg RNA was reverse transcribed to cDNA with the High-Capacity RNA-to-cDNA Kit (Thermo Scientific 4387406). PCR was performed using GoTaq Green Master Mix (Promega M7122) with the following primers: F: 5’-CCCCTCCTCTCCATACTCCT-3’; R: 5’-CCTCTACGTGCTCAGGATCC-3’. PCR products were then run in a 2% agarose gel.

### Western blot and nucleus-cytoplasmic fractionation

HEK293 cells were lysed in cold RIPA buffer with protease/phosphatase inhibitor cocktail. Lysates were incubated on ice for 15min at 4°C with vortex every 5min. Lysates were then centrifugated at 10,000rpm for 10min (4°C), and the supernatant transferred to a new tube. Samples were prepared for WB by diluting 1:1 with 2X Laemmli buffer with 5% β-mercaptoethanol and incubating for 5min at 95°C. Samples were run in a 4-20% polyacrylamide gel using the Mini-PROTEAN electrophoresis Bio-Rad system. Proteins were transferred to a PVDF membrane using the Trans-Blot turbo Bio-Rad system. Following transfer, the membrane was blocked for 1h with 5% BSA in TBS with 2% Tween-20 (TBST) followed by overnight incubation with primary antibody at 4°C. After 4x 10-min washes with TBST, the membrane was incubated with secondary antibody for 1–2 h at room temperature and washed again 4 times with TBST. Horseradish peroxidase (HRP) was detected using SuperSignal West Dura kit, Thermo Scientific. Western blot images were acquired by Azure 600 (Azure Biosystems) and quantified by an ImageJ software package. Primary antibodies: mouse anti-Flag (1:1,000, Sigma F1804), mouse anti-GAPDH 6C5 (1:10,000, Abcam ab8245), rabbit anti-H3 (1:5,000, Abcam ab1791). Secondary antibodies: goat anti-rabbit HRP (1:5,000; Jackson ImmunoResearch Labs 111-036-047), goat anti-mouse HRP (1:5,000; Jackson ImmunoResearch Labs 115-036-072).

For the fractionation experiment, the cells were washed with PBS, collected in 1ml cold PBS, and centrifugated at 100G for 5min (4°C) to pellet the cells. The supernatant was removed, and the cells were resuspended by vortexing in 200μl cold NP40 buffer (150mM NaCl, 50mM HEPES pH: 7.4, 1% NP40). The suspension was incubated for 10min on ice and centrifuged for 10min at 7000G to pellet nuclei and cell debris. The supernatant was transferred to a new tube (cytosolic fraction) and prepared for WB by diluting 1:1 with 2X Laemmli buffer with 5% β-mercaptoethanol and incubating for 5min at 95°C. The pellet was resuspended by vortexing in 200 μl cold NP40 buffer containing 1U/ml Benzonase. The samples were incubated in a rotator at 4°C for 1hr and then centrifuged for 10min at 7000G. The supernatant was transferred to a new tube (nuclear fraction) and prepared for WB by diluting 1:1 with 2X Laemmli buffer with 5% β-mercaptoethanol and incubating for 5min at 95°C.

### Immunofluorescence

HEK293 cells were cultured in PDL-coated (100μg/ml) glass coverslips. Two days after treatment with ASOs, cells were subjected to immunofluorescence staining. Briefly, cells were washed one time with PBS, then fixed for 10min in 4% cold PFA. After 3x PBS washes, cells were permeabilized with 0.1% Triton in PBS for 10min, blocked with 5% BSA for 1hr, and incubated for 1hr in first antibody diluted in blocking buffer. Next, cells were washed 3 times with PBS and incubated for 1hr in secondary antibody. Finally, the nucleus was stained using the NucBlue reagent (Hoechst 33342) following the manufacture instructions. The coverslips were mounted into microscopic slides using Prolog Glass mounting media, and images were taken using the EVOS-FL microscope. Primary antibodies: mouse anti-Flag (1:500, Sigma F1804), rabbit anti-β-tubulin (1:500, Abcam ab6046). Secondary antibodies: goat anti-mouse Alexa Fluor 488 (1:1,000, Thermo Scientific A-11001), goat anti-rabbit 555 (1:1,000, Thermo Scientific A-31572).

### Luciferase assay

The promoter of bovine RHO was cloned into a pGL3 luciferase reporter vector (Promega E1741). HEK293 cells were transfected with plasmids expressing the RHO reporter vector, CRX-HA, NRL-HA, eGFP, and either Flag-NR2E3^WT^ or Flag-NR2E3^c.932G>A^ using FuGene HD transfection reagent. ASOs (100nM) were transfected 24hrs after the plasmid’s transfection. Luciferase activity was performed 48 hrs following ASO transfection with the Firefly Luciferase assay system (Promega E4030), following the manufacturer instructions. Both luciferase activity and eGFP fluorescence were read using a Tecan Spark 20M microplate reader.

